# Shape Analysis of Coronary Flow Waveforms using Singular Value Decomposition

**DOI:** 10.64898/2026.08.26.743980

**Authors:** Victoria E. Sturgess, Noah A Schenk, Joshua W Ziegele, Salman I Essajee, Johnathan D Tune, Indika Rajapakse, C. Alberto Figueroa, Daniel A. Beard

## Abstract

Coronary flow waveforms have a distinct diastolic-dominant shape with periods of low or retrograde flow during systole. While the general waveform shape has been attributed to complex interactions between cardiac and vascular mechanics, there is limited research into the variability in coronary flow waveforms and what this variability may reveal about cardiac function.

This work presents a shape analysis of left anterior descending artery (LAD) flow waveforms using Fourier transforms and Singular Value Decomposition (SVD) performed on baseline data collected from 32 pigs. Pigs included in the study reflect two breeds (Ossabaw and Yorkshire) and three different experimental conditions (lean-control, lean-paced, and obese-paced). Fourier transforms were used to decompose the waveforms into 15 harmonics for each pig. An SVD analysis is then used to extract temporal patterns of the waveforms. Correlations between pig-specific coefficients for the SVD modes and clinical metrics were used to investigate physiological explanations of LAD waveform variability.

Temporal LAD flow patterns of the second SVD mode are significantly correlated with heart rate. The third SVD mode significantly correlates with mean blood pressure and maximum hyperemic flow. Furthermore, the fourth SVD mode is weakly correlated with left-ventricular end diastolic pressure and endocardial-epicardial flow ratios.

This work demonstrates that LAD flow waveforms can be broken down into temporal patterns that correlate with physiological features. Furthermore, this shape-analysis method allows for waveform reconstruction and simplifies visualization of the temporal patterns identified using SVD, an advantage over existing methods that focus on characterizing flow waveforms by points of interest.

## 1 Introduction

The shapes of coronary arterial flow waveforms result from complex interactions between hemodynamics and cardiac muscle mechanics. During early systole, cardiac muscle contracts and exerts pressure on the outside of myocardial blood vessels. This extravascular pressure is heterogeneous across the heart wall, with the largest values near the ventricular cavity (endocardium) and the smallest at the outer surface of the heart (epicardium) [1, 2]. The spatial and temporal variations of extravascular pressure in the left ventricle create unique hemodynamic patterns, including diastolic-dominant arterial flow, periods of low or reversed arterial flow during systole, and different flow patterns between endocardial and epicardial vessels [3].

Despite these complex temporal patterns, quantitative metrics of coronary hemodynamics often rely on simple quantities such as time-averaged flow over a cardiac cycle. Endocardial/epicardial flow ratios (Endo-Epi ratio) provide an estimate of time-averaged regional flow differences. These ratios can be measured using radiotracers and positron emission tomography [4] or through the perfusion of labeled microspheres [5]. Another metric, coronary flow reserve (CFR), measures the ratio of average flow in a hyperemic state compared to a baseline state. While these averaged quantities provide important functional information, we hypothesize that additional structural and functional information may be encoded within the coronary flow waveforms.

To date, there exists limited work focused on analyzing temporal patterns of coronary flow waveforms. Qualitative descriptions have included classification based on the presence or absence of reverse arterial flow during mid to late systole [6]. Wave intensity profiles provide a more quantitative assessment of coronary velocity and pressure waveforms by identifying accelerating and decelerating waves from the proximal and distal ends of coronary arteries. Studies using this method suggest that velocity accelerations in early diastole are the result of backward traveling “suction waves” caused by ventricular relaxation [7, 8]. Recently, Geddes and Randles identified a minimal number of points of interest required to characterize the shape of coronary velocity waveforms and used this technique to study the shape of velocity waveforms in synthetically generated datasets [9, 10]. However, we submit that more work is required to understand how the shapes of *in-vivo* coronary flow waveforms differ between subjects and how these differences are related to underlying hemodynamics.

Fourier transforms and singular value decomposition (SVD) have been used to analyze periodic gait patterns. Extensive publications in this area have provided correlations between walking patterns and individual characteristics such as height, weight, walking speed, and gender [11, 12]. We hypothesize that similar techniques can be applied to study cardiovascular hemodynamic patterns, specifically coronary arterial flow waveforms.

In this work, we characterize left anterior descending artery (LAD) coronary flow waveforms using Fourier transforms and SVD. We perform a quantitative shape analysis assessment of these waveforms using Fourier modes and SVD analysis to identify temporal patterns within the dataset. Correlations between SVD temporal patterns and experimental data, such as heart rate and blood pressure, are unveiled.

## 2 Methods

### 2.1 Experimental Data

LAD flow waveforms, left ventricular pressure, aortic pressure, and other experimental data were collected from anesthetized swine at the University of North Texas Health Science Center. All protocols were approved by the appropriate Institutional Animal Care and Use Committees in accordance with the Guide for the Care and Use of Laboratory Animals (NIH Pub. No. 85–23, Revised 2011). In total, recordings from 32 pigs (24 Ossabaw pigs and 8 Yorkshire Domestic pigs) were used within this study. The pigs were subdivided into five groups based on the breed and experiment type: lean control Ossabaw (10), lean paced Ossabaw (8), obese paced Ossabaw (6), lean control Yorkshire (7), and lean paced Yorkshire (1). Paced animals underwent a right-ventricular 180 beats-per-minute pacing protocol for 4 weeks. Pacemakers were turned off for these animals before data collection. Lean animals were fed a standard diet, while obese animals were fed an excess-calorie, high-fat, high-fructose diet. Details on the pacing protocol, diet, and surgical protocol are described in Tune et al. [13]. Data included in this study represent baseline states, stabilization of hemodynamic measurements following instrumentation, and were recorded as part of other experiments (Tune et al. [13] and Essajee et al. [14]).

Quantitative metrics were used to describe the hemodynamic, metabolic, and physiological states of the pigs in the previous studies. Body weights and heart weights (HW) were collected before and after the experiment, respectively. For each pig, hemodynamic measurements of left ventricular pressure, aortic pressure, and LAD coronary flow were collected with sampling frequencies of 1000 Hz. For thsi study, the derivative of left ventricular pressure with respect to time (dPLV/dt) is calculated. Maximal values of dPLV/dt (maximal contractility during systole) are used to define the start and end of each cardiac cycle and calculate heart rate (HR). Hemodynamic measurements are collected for 30 cardiac cycles and are used to calculate maximum and minimum left ventricular pressure (PLV max and min), maximal and minimum dPLV/dt, and mean blood pressure (MBP). Additionally, left ventricular end diastolic pressure (LVEDP) is defined as the left ventricular pressure during contraction when dPLV/dt is 20% of its maximal value, and the average value of LVEDP is recorded for the 30 cardiac cycles. Flow-based metrics collected from this baseline period include average flow across the 30 cardiac cycles (Avg Flow) and average flow per gram of heart weight (HW-Norm Avg Flow). A flow pulsatility metric is defined as the difference between the maximal and minimal blood flow during a cardiac cycle.

Variability of cardiac cycle duration is also calculated. Cardiac cycle variability (CCV) for *N*_*i*_ cardiac cycles is defined for each pig as the sum of the absolute values of the differences between each cardiac cycle duration *d*_*i*_ and the pig’s average cardiac cycle duration *d*_*avg*_. This metric is calculated as a unit of time (milliseconds) and as a percentage of the average cardiac cycle duration, see equations 1 and 2, respectively.

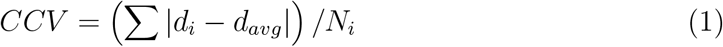

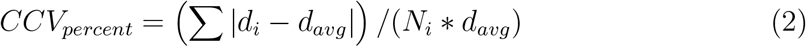

In the previous experiments, the following additional information was collected for some of the pigs: blood oxygen measurements (21/32 pigs), endo-epi regional blood flow estimates (21/32 pigs), and coronary stress tests (12/32 pigs). Oxygen extraction was determined from arterial and coronary venous blood samples. Injections of neutron-activated microspheres were used to estimate endo-epi flow ratios. Coronary stress tests were performed by administering regadenoson after the baseline period. The average flow of each cardiac cycle is calculated and is used to determine a baseline flow prior to administration of regadenoson and a maximal hyperemic flow. A CFR value is calculated as the ratio of maximal hyperemic flow to the pre-regadenoson baseline flow (equation 3). The change in flow in response to regadenoson (Δ Regadenoson) is calculated as the difference between hyperemic flow and pre-regadenoson baseline flow (equation 4).

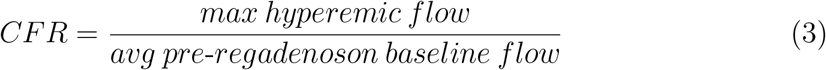

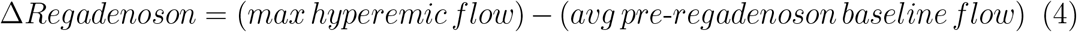

Further analysis on the coronary flow waveforms from the baseline 30 cardiac cycles of each pig is described in the following sections: Fourier Feature Extraction, Singular Value Decomposition, and Physiological Interpretation of SVD Modes. Figure 1 depicts these steps in an overview of the methodology.

**Figure 1:**
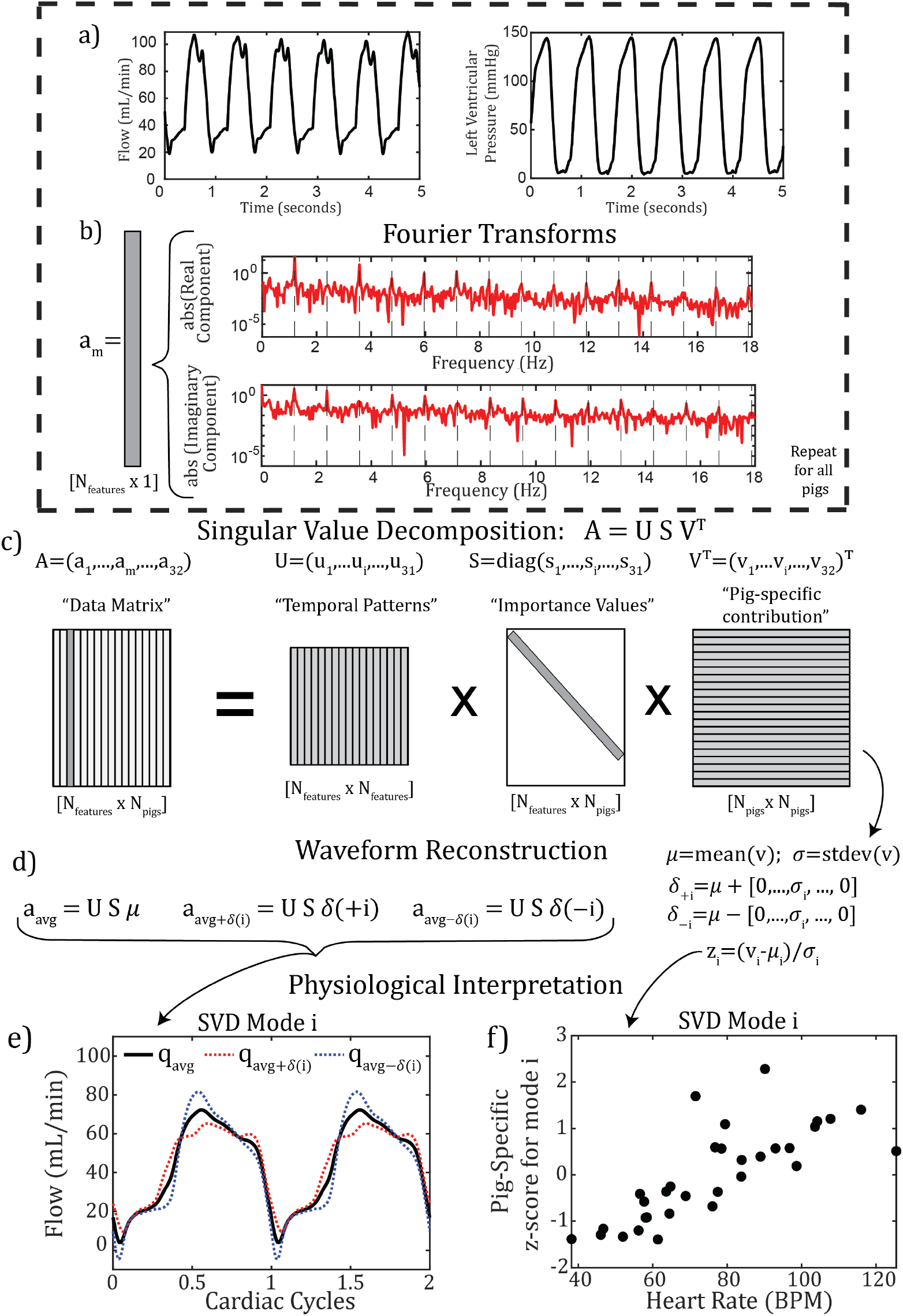
The various stages included within the computational workflow: (a) collection of hemodynamic data; (b) collection of information in the Fourier transform stored at harmonics of the cardiac cycle frequency (reflected by the dashed lines) and stored in vector *a*_*m*_; (c) singular value decomposition of the *A* data matrix; (d) construction of new vectors representing the average and the average plus or minus a standard deviation for SVD mode *i*; (e) visualization of these trends for the i^th^ SVD mode; and (f) visualization of trends between heart rate and the z-score values of the right singular vector of SVD mode i.

### 2.2 Fourier Feature Extraction

Discrete Fourier transforms (DFTs) are performed on the coronary flow waveform for each pig. From each DFT, complex values representing Fourier modes at harmonics of the cardiac cycle frequency are calculated.

To determine whether the DFTs sufficiently capture the complexity of the original data, new waveforms are reconstructed using sinusoidal functions at harmonics of the cardiac cycle frequency (equation 5), where the variable *f* represents the individual pig’s cardiac cycle frequency, *N*_*k*_ is the number of harmonics, and *t* is time. Variables 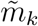 and 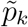 represent the magnitude and phase of the *k*^*th*^ harmonic.

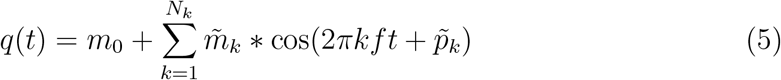

The zero-frequency harmonic of the raw coronary flow waveform, *m*_0_, equivalent to the time-averaged coronary flow, is not included in the number of harmonics represented by *N*_*k*_. For each pig, *q*(*t*) waveforms are reconstructed with *N*_*k*_ values ranging from 1 to 50 and a sampling frequency of 1000 Hz to match the original data.

Information about the magnitude and phase of each harmonic is also represented as a complex number: (*α*_*k*_ + *β*_*k*_*i*). In this Cartesian format, the magnitude of the harmonic is given by Equation 6 and the phase of the harmonic is given by Equation 7, where tan^−1^ returns the four-quadrant inverse tangent. Cartesian formats with the real and imaginary components of the complex number are used in the subsequent SVD analysis to avoid issues that would arise from the periodicity of phase angles (i.e., 0 and 2*π* radians being equivalent).

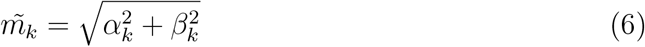

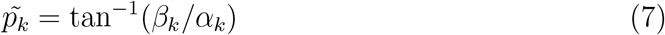

For each pig, the reconstructed waveforms are compared to the original flow waveform using mean squared error (MSE) in equation 8, where *Q*(*t*) is the 30 cardiac cycles of the raw flow waveform, *q*(*t*) is 30 cardiac cycles of the periodic reconstructed raw waveform, *T* is the number of time points. A normalized mean square error or NMSE is calculated by dividing by the temporal standard deviation of the raw coronary flow (equation 9).

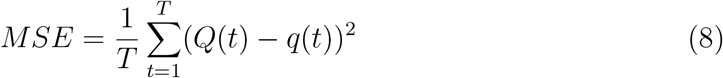

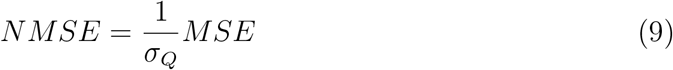

To determine the number of Fourier modes required to capture the coronary flow waveform, MSE and NMSE values are calculated for values of *N*_*k*_ up to 50. For subsequent SVD analyses, only the first 15 harmonics are used, *N*_*k*_ = 15.

### 2.3 Singular Value Decomposition

For a given pig *m*, a 31x1 vector *a*_*m*_ is constructed as a low-dimensional approximation of its LAD coronary flow waveform using the magnitude of the 0-frequency harmonics and the real and imaginary DFT components of the first 15 harmonics of the pigs cardiac cycle frequency. After assembling *a*_*m*_ vectors for all 32 pigs, they are combined into a data matrix (*A*) with size *N*_*features*_ × *N*_*pigs*_ or 31 × 32.

SVD is performed to decompose matrix *A* into three matrices of left singular vectors (*U*), right singular vectors (*V*), and singular values (*S*), shown in Figure 1d.

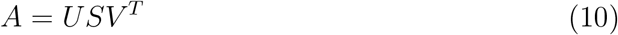

The *U* and *V* matrices have a size of *N*_*features*_ × *N*_*features*_ and *N*_*pigs*_ × *N*_*pigs*_, respectively. The rows of *U* and *V* ^*T*^ represent the left and right singular vectors, respectively. The *S* matrix has the same size as the data matrix *A* (*N*_*features*_ × *N*_*pigs*_). Matrix *S* contains positive values only along its diagonal and zeros elsewhere. The diagonal entries, referred to as singular values, are organized in decreasing order.

Together, the i^th^ left singular vector, i^th^right singular vector, and i^th^ singular value are referred to as the i^th^ SVD mode. Given the structure of matrix *A*, the left singular vectors provide information about the temporal patterns associated with each mode, and the right singular vectors provide information about the contribution of that mode for each pig.

Singular values contain information about the importance of each mode in comparison to the other modes. The percent of variance explained for the i^th^ SVD mode is defined by the square of the i^th^ singular value (*s*_*i*_) divided by the summation of the square of all singular values. Cumulative variance for the i^th^ SVD mode is defined as the summation of the percent explained variance for all SVD modes up to and including the i^th^ SVD mode:

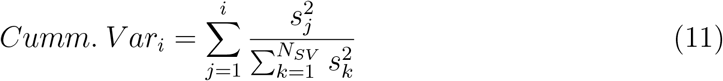

Additionally, the average and standard deviation of the i^th^ right singular vector (*v*_*i*_) are denoted as *µ*_*i*_ and *σ*_*i*_, respectively.

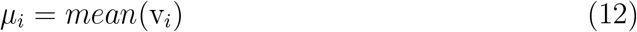

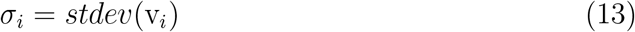

Right singular vectors are converted to vectors of z-scores using the average and standard deviations for the given right singular vector. The vector (*z*_*i*_) denotes the pig-specific z-scores of the i^th^ right singular vector and provide information about the contribution of the i^th^ mode for each pig compared to the mode average *µ*_*i*_.

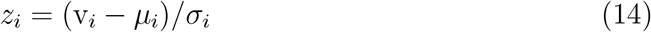

Vectors *µ* and *σ* are the row average and row standard deviation of matrix *V* ^*T*^ and contain the values *µ*_*i*_ and *σ*_*i*_ for each right singular vector.

### 2.4 Waveform Reconstruction

The column vector *a*_*m*_ could be reconstructed as a linear combination of the information in the SVD modes where *u*_*i*_ is the i^th^ left singular vector, *s*_*i*_ is the i^th^ singular value, v_*i*_(*m*) is the value of the m^th^ entry of the i^th^ right singular vector, and *N*_*SV*_ is the number of SVD modes. Equivalently, the vector *a*_*m*_ could be defined through matrix-vector multiplication where vector 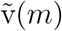 is the m^th^ column of *V* ^*T*^, containing the m^th^ value of each right singular vector.

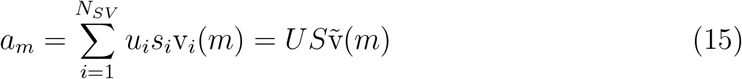

Using similar methods, a representative set of features (*a*_*avg*_) is constructed for the average coronary waveform by replacing v_*i,j*_ with *µ*_*i*_ from equation 12. This is equivalent to matrix-vector multiplication using matrices *U* and *S* and vector *µ*.

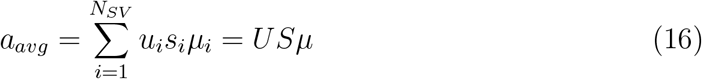

Additionally, for each SVD mode, two new sets of features *a*_*avg*+*δ*(*i*)_ and *a*_*avg*−*δ*(*i*)_ are constructed to represent average features plus or minus one standard deviation of the i^th^ SVD mode. These representative vectors are modified from *a*_*avg*_ by adding or subtracting the product of the i^th^ left singular vector, singular value, and *σ*_*i*_ or the standard deviation of the i^th^ right singular vector from equation 13. Matrix-vector multiplication could also be used to define *a*_*avg*+*δ*(*i*)_ and *a*_*avg*−*δ*(*i*)_. Vectors *δ*(+*i*) and *δ*(−*i*) are defined as *µ ±* [0, …, *σ*_*i*_, …, 0], where only the i^th^ value of the *δ*(+*i*) and *δ*(−*i*) vectors differ from the *µ* vector.

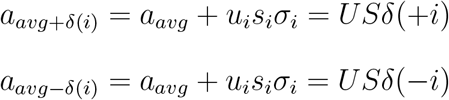

These representative sets of features are converted into representative flow waveforms (*q*_*avg*_, *q*_*avg*+*δ*(*i*)_, and *q*_*avg*−*δ*(*i*)_) with a process similar to equation 5.

### 2.5 Physiological Interpretation of SVD modes

The contribution of each SVD mode to the shape of the coronary flow waveform are visualized by plotting representative waveforms (*q*_*avg*+*δ*(*i*)_ and *q*_*avg*−*δ*(*i*)_) against the average coronary waveform(*q*_*avg*_). These graphs display temporal patterns associated with higher or lower contributions of the i^th^ SVD mode (Figure 1f).

Additional physiological interpretation for each SVD mode is determined using linear regressions between values of the z-score vectors (*z*_*i*_) and data metrics described in the *Experimental Data* section of the methods. Pearson coefficients (r), coefficient of determination (R^2^), and p-values are calculated for the linear regressions. A Bonferroni adjustment is used to control for multiple comparisons and reduce the likelihood of false positives. The top five modes are each compared to 20 experimental metrics. Therefore, an adjusted p-value of 0.05*/*(20 × 5) = 0.0005 is used as a threshold for significance in correlations between SVD modes and experimental metrics. Scatter plots are used to visualize the correlation between z-scores of SVD modes and data metrics (Figure 1g).

Estimates of hyperemic flow are constructed with single- and multi-variable linear regression models using SVD modes and average baseline flow. The quality of these regression models compared to measured hyperemic flow is evaluated using MSE and R^2^ values.

### 2.6 Representative Waveforms for Experimental Groups

A comparison between Ossabaw pigs from the three experimental conditions (Lean Control, Lean Paced, and Obese Paced) is performed by constructing a representative set of features and a waveform for each condition. Experimental group means of right singular vectors *µ*(*LeanControl*), *µ*(*LeanPaced*), and *µ*(*ObesePaced*) are constructed, where i^th^ entry of each *µ*(*X*) vector represents the mean all entries corresponding to Ossabaw pigs in experimental group X of the i^th^ right singular vector. Average sets of features (*a*_*avg*_(*LeanControl*), *a*_*avg*_(*LeanPaced*), and *a*_*avg*_(*ObesePaced*)) for each experimental group are constructed using equation 16 and replacing *µ* with the respective experimental group means *µ*(*X*). Representative waveforms for each experimental group are constructed with a process similar to equation 5. Experimental group means of z-values *z*_*i*_(*LeanControl*), *z*_*i*_(*LeanPaced*), and *z*_*i*_(*ObesePaced*) are also calculated for each SVD mode i.

## 3 Results

### 3.1 Fourier Features

DFTs were used to convert the coronary flow waveforms from the temporal domain to the frequency domain. Analysis of the frequency space showed that most occurred at the lower harmonics of the cardiac cycle frequency (Figure 1c). Additionally, low-frequency spikes, potentially associated with breathing artifacts, were observed for some pigs.

To determine the number of harmonics or Fourier modes required to capture the coronary flow waveform, *N*_*k*_, reconstructed waveforms were compared to the original waveforms. Trends of MSE and NMSE for *N*_*k*_ values ranging from 1 to 50 are displayed in Figure 2. After 10-15 Fourier modes, the MSE and NMSE became relatively constant, and information provided by additional modes did not improve the accuracy of the reconstructed waveforms. Therefore, a threshold of 15 Fourier modes was chosen to represent the data for the remaining analysis. A sensitivity analysis of results presented in the following SVD sections was performed with *N*_*k*_=10 and 20, and similar overall results were observed.

**Figure 2:**
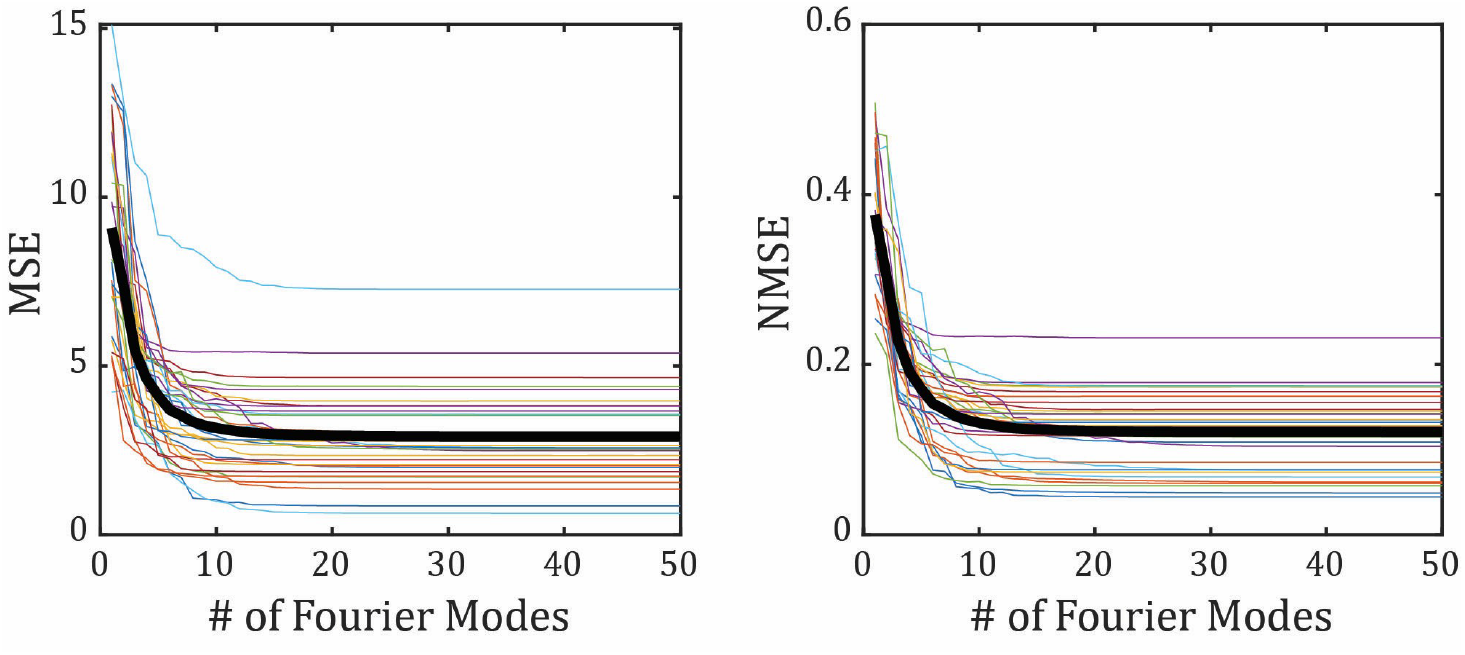
Impact of the number of Fourier modes (*N*_*k*_) on the MSE (left) and NMSE (right) between the original and reconstructed coronary flow waveforms. Trends for each pig are shown by the thin colored lines, and the thick black line represents the average MSE or NMSE across all pigs.

Examples of three coronary flow waveforms and their reconstructed signals for low, moderate, and high CCV and NMSE are shown in Figure 3 a-c. In general, pigs with low CCV (reflecting variability of cardiac cycle duration) had lower NMSE values, and this trend remained whether CCV was calculated as a time or a percent of the average cardiac cycle duration (Figure 3 d and e). Some pigs had coronary flow waveforms that changed shapes over the 30 cardiac cycles in which data was collected (Figure 3b), resulting in moderate NMSE. For pigs with high CCV, cardiac cycles oscillated between shorter and longer durations and often caused high NMSE values. For example, in Figure 3c, the high-frequency flow dip that occurred in early systole was not accounted for in the reconstructed waveforms. The frequency versus magnitude trends (example shown in Figure 1c) appeared to show spectral leakage for pigs with high CCV compared to pigs with low CCV.

**Figure 3:**
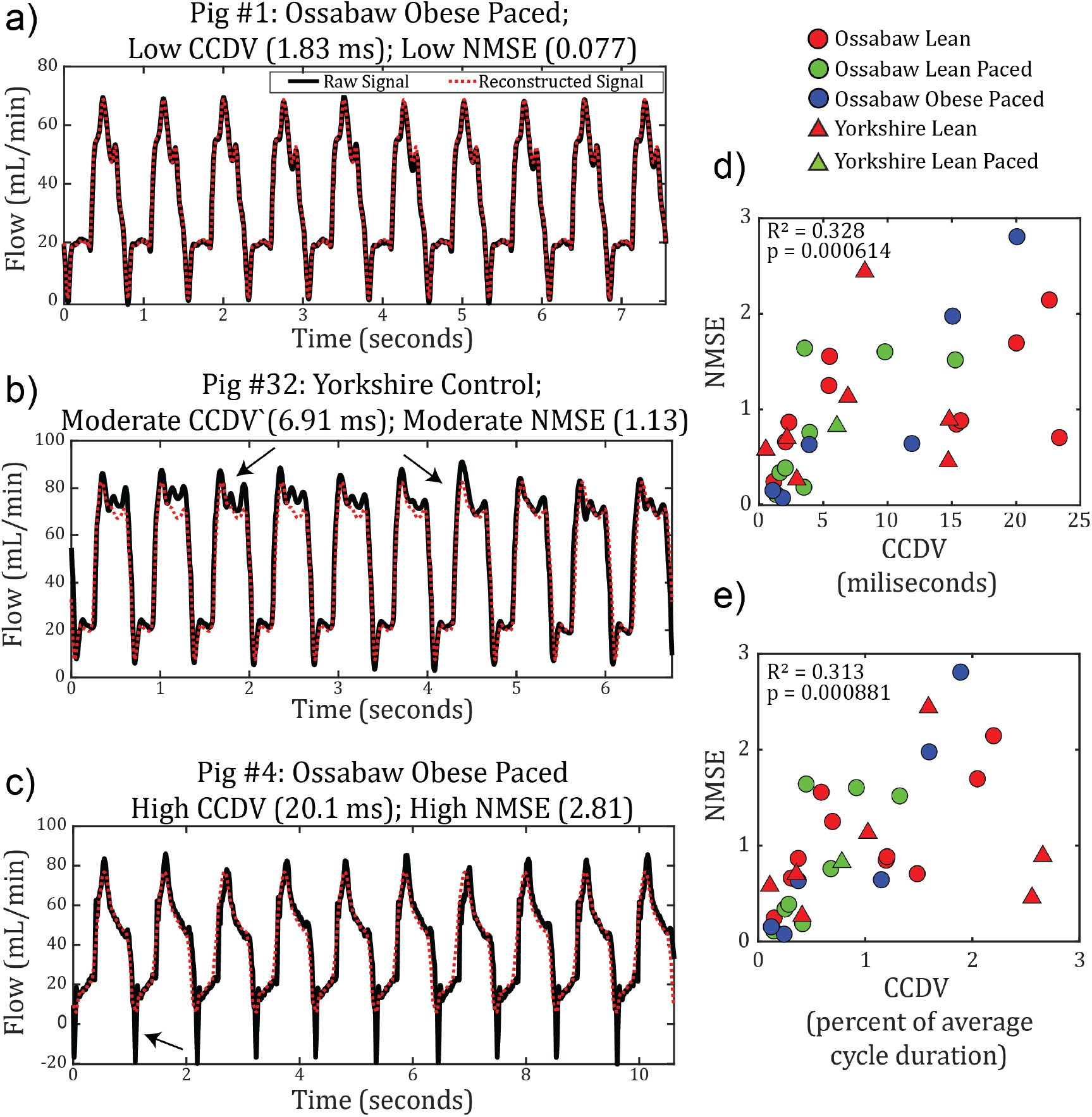
Examples of the raw (black lines) and reconstructed flow waveforms (red dashed lines) for three pigs (left) and the relationship between CCV and NMSE (right). The low CCV and low MSE pigs (a) had good agreement between the raw and reconstructed waveforms. (b) Illustrates a raw waveform with shape changes throughout the 30 cardiac cycles (shown by arrows), causing high MSE. The reconstructed signal in example (c) does not capture the sharp dip in early systole (shown by the arrow), leading to high NMSE. The relationship between CCV and NMSE is reported with CCV in milliseconds (d) and as a percentage of the cardiac cycle (e).

### 3.2 Singular Value Decomposition

The *N*_*feature*_ × *N*_*pigs*_ data matrix was assembled with information on the 31 features (zero-frequency magnitude and the real and imaginary components of the first 15 harmonics of the cardiac cycle frequency) for the 32 pigs. Decomposition of this data matrix into three matrices of left singular vectors, singular values, and right singular vectors was performed using SVD. The first SVD mode accounted for 97% of the variance, and the top five SVD modes accounted for 99.7% of the variance (Figure 4a, red dot). Modal contributions, shown in Figure 4b, were calculated as *s*_*i*_ × (*µ*_*i*_ *± σ*_*i*_), where *s*_*i*_ was the i^th^ singular value and *µ*_*i*_ and *σ*_*i*_ were the mean and standard deviation of the i^th^ right singular vector, respectively. The contribution of the first SVD mode was negative for all pigs. For all other SVD modes, the mean modal contributions were approximately zero, and the range of nodal contributions included both positive and negative values. With higher SVD modes, the singular value (*s*_*i*_) decreased, and the corresponding variability of the modal contributions also decreased.

**Figure 4:**
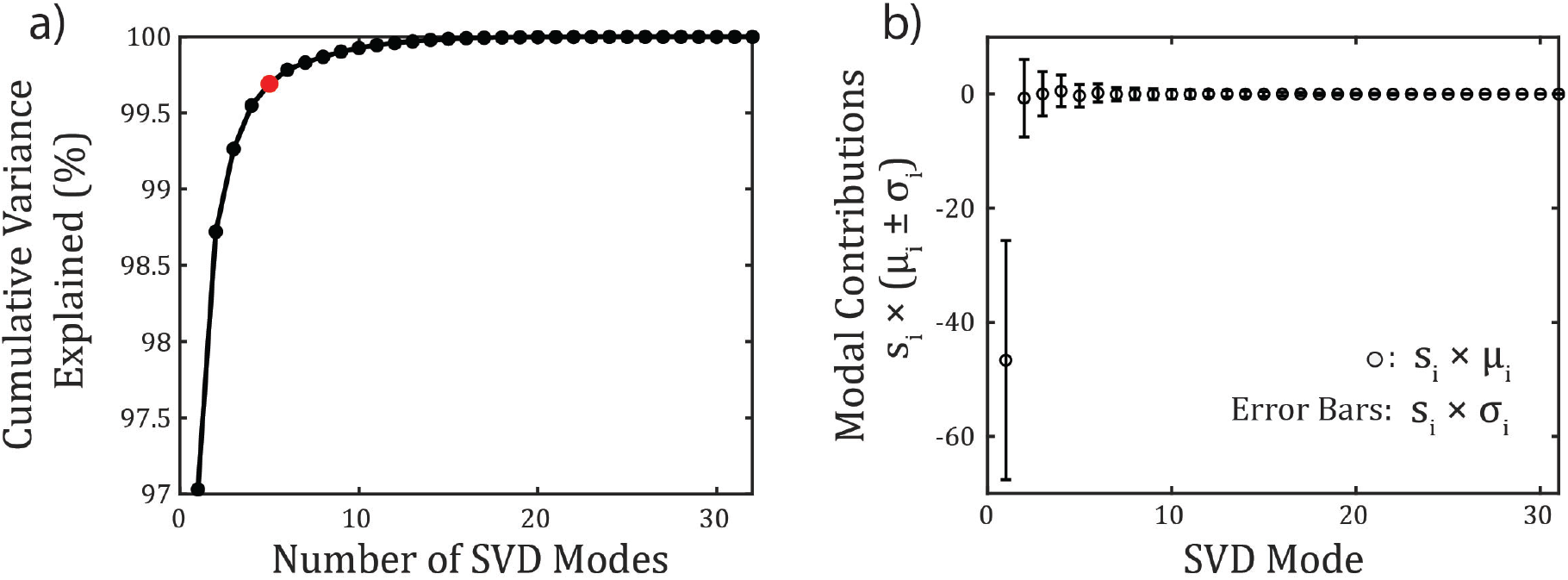
Cumulative variance (a) and modal coefficients (b) from the SVD analysis. For the modal coefficients, open circles represent the products of the singular values (*s*_*i*_) and the mean values of the right singular vectors (*µ*_*i*_). The heights of the error bars represent the product of singular values and the standard deviation of the right singular vectors (*σ*_*i*_).

### 3.3 Physiological Interpretation of Singular Vectors

To gain physiological insight for the top five SVD modes, representative waveforms for the average were compared to the average plus/minus one standard deviation of mode i (Figure 5). Additionally, correlations between the z-scores of the right singular vectors and experimental data were assessed. P-values and *R*^2^-values for all correlations between the top 5 SVD modes and 20 experimental variables are displayed in Figure 6.

**Figure 5:**
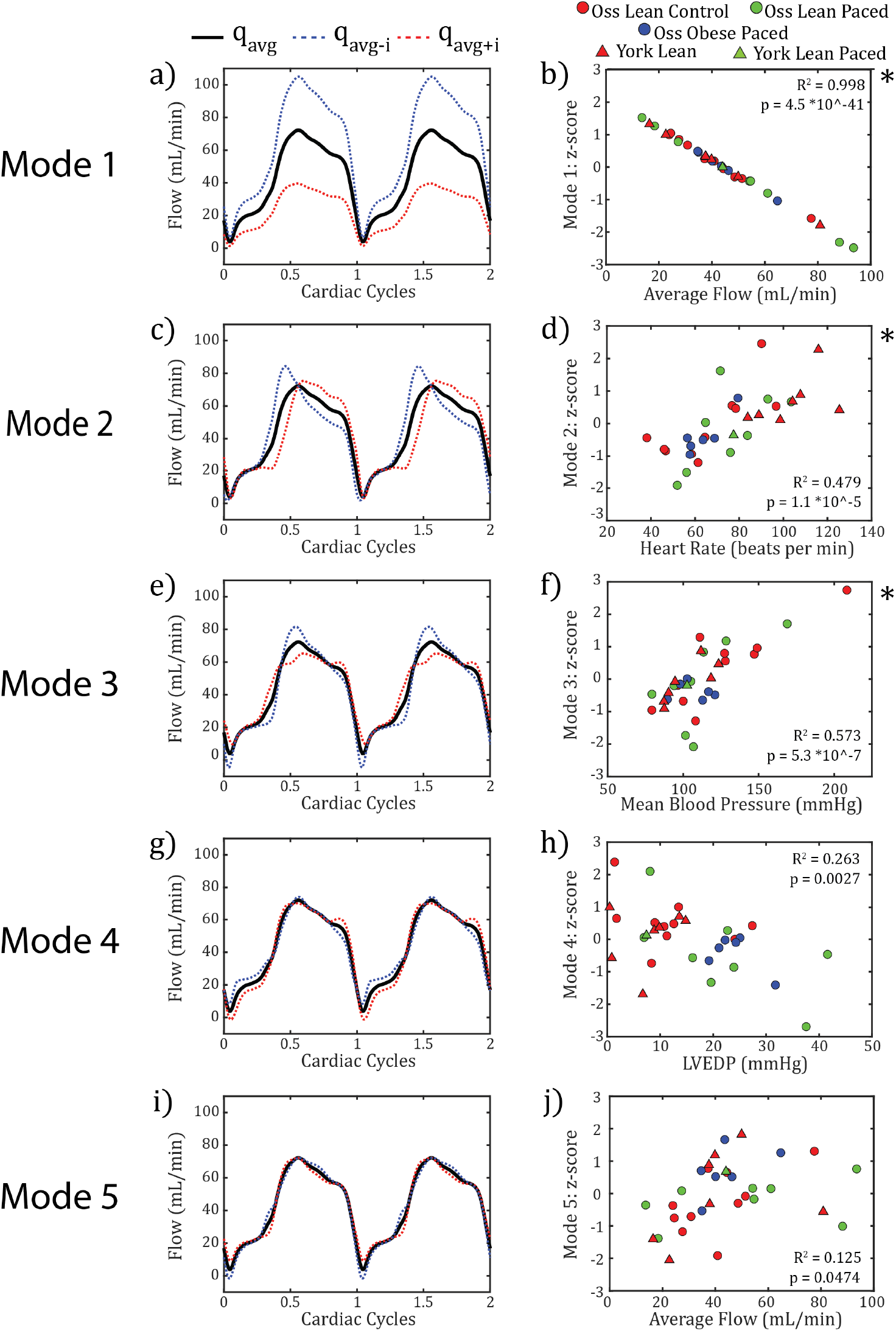
Temporal trends of the first five SVD modes (left) and correlations between z-scores of right singular vectors to clinical data (right). Temporal trends include the a waveform representing the average pig *q*_*avg*_ in black, the average plus one standard deviation of the i^th^ SVD mode *q*_*avg*+*δ*(*i*)_ in red dashes, and the average minus one standard deviation of the i^th^ SVD mode *q*_*avg*−*δ*(*i*)_ in blue dashes. All waveforms begin in early systole and include two complete cardiac cycles. A star to the right of the correlation graph indicates a p-value less than the adjusted alpha value of 0.0005.

**Figure 6:**
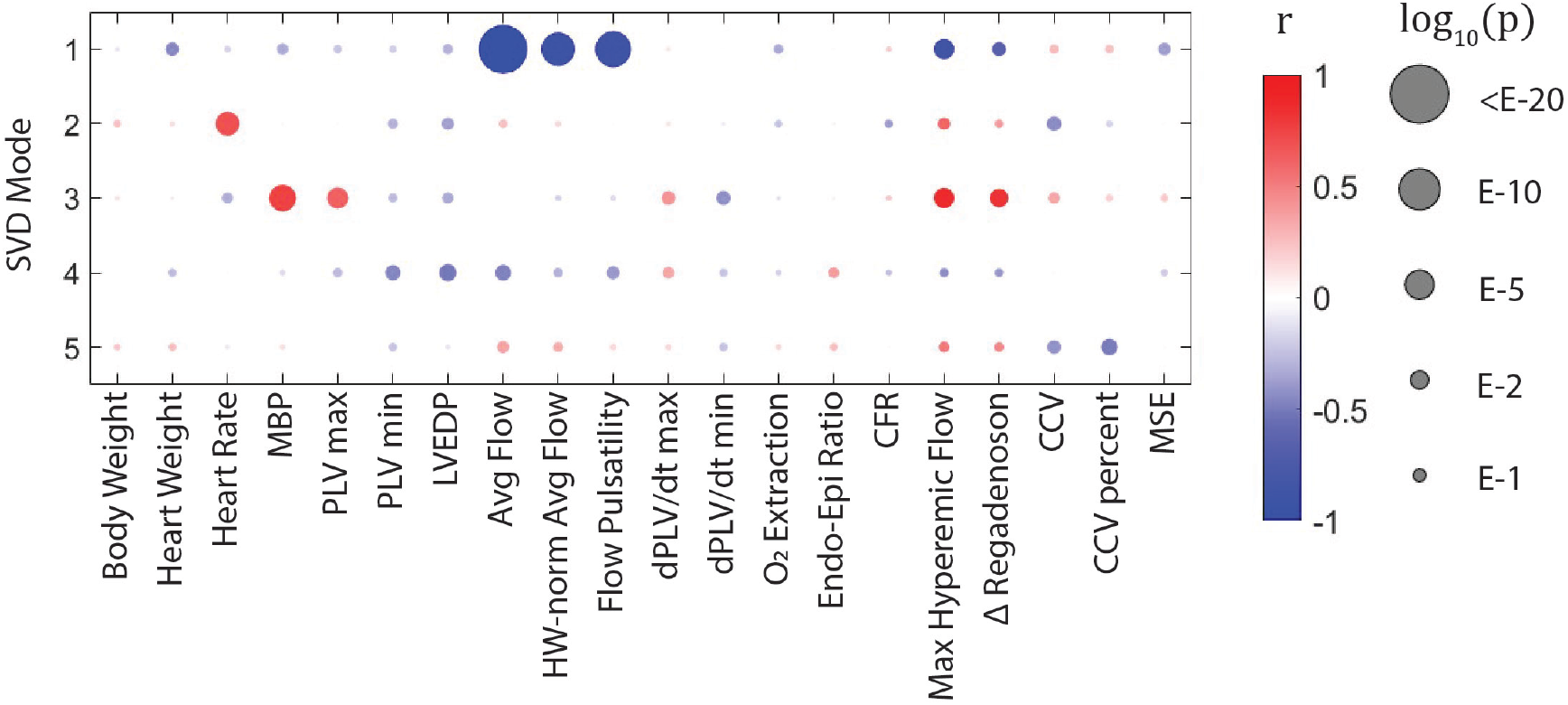
Visualization of the correlation analysis between SVD modes and experimental data metrics for the raw waveform analysis.

The first SVD mode affected the magnitude of the waveform. The average plus one standard deviation (*q*_*avg*+*δ*(1)_) decreased the systolic and diastolic portions of the coronary flow waveform (Figure 5a). Opposite trends were observed with the average minus one standard deviation (*q*_*avg*−*δ*(1)_). Therefore, a near-perfect negative correlation was observed between the first SVD mode and the average flow (Figure 5b). This SVD mode also had significant negative correlations with average flow per gram heart weight, flow pulsatility, and maximal hyperemic flow (Figure 6).

The second SVD mode affected the phase of the waveform. Larger contributions of the second mode delayed the diastolic peak and appeared to increase the length of systole (Figure 5c). Furthermore, this mode also altered the relative contribution of early and late diastolic flow. Decreases in the second mode increased flow in early diastole and decreased flow in late diastole compared to the average flow profile. A significant positive correlation was found between the second mode and heart rate (Figure 5d).

The third SVD mode affected the slope of the early diastolic rise in coronary blood flow (Figure 5e). Increasing contributions of this mode caused a more gradual change in slope, whereas lower contributions caused a more rapid slope. This mode also altered the magnitude of both the early systolic flow minimum and the early diastolic flow maximum, with lower contributions of this mode causing a lower minimal flow and higher maximal flow. Mean blood pressure showed a significant positive correlation with the third SVD mode (Figure 5f).

The fourth SVD mode affected the systolic portion of the coronary flow waveform, with higher modal contributions causing higher flow during systole (Figure 5g). In addition, a late diastolic bump appeared in the coronary flow waveform with higher contributions of this mode. The fourth SVD mode had no significant correlations with any experimental data. However, a non-significant negative correlation was observed between LVEDP and the fourth SVD mode (Figure 5h).

The fifth SVD mode affected the early systole dip in the coronary flow waveform and had no significant correlations with the data metrics.

No significant correlations with any of the top 5 SVD metrics were observed for CFR or the Endo-Epi ratio metrics (Figure 6). Within the top five SVD modes, the Endo-Epi ratio metric had a non-significant correlation (p=0.0873) with the fourth SVD mode (Figure S1).

Significant correlations between flow rates (average baseline flow and maximal hyperemic flow) and the first and third SVD modes were observed, and corresponding scatter plots are shown in Figure 7. While SVD mode 1 had a significant correlation with both flow measurements, the third SVD mode only showed a significant correlation with maximal hyperemic blood flow.

**Figure 7:**
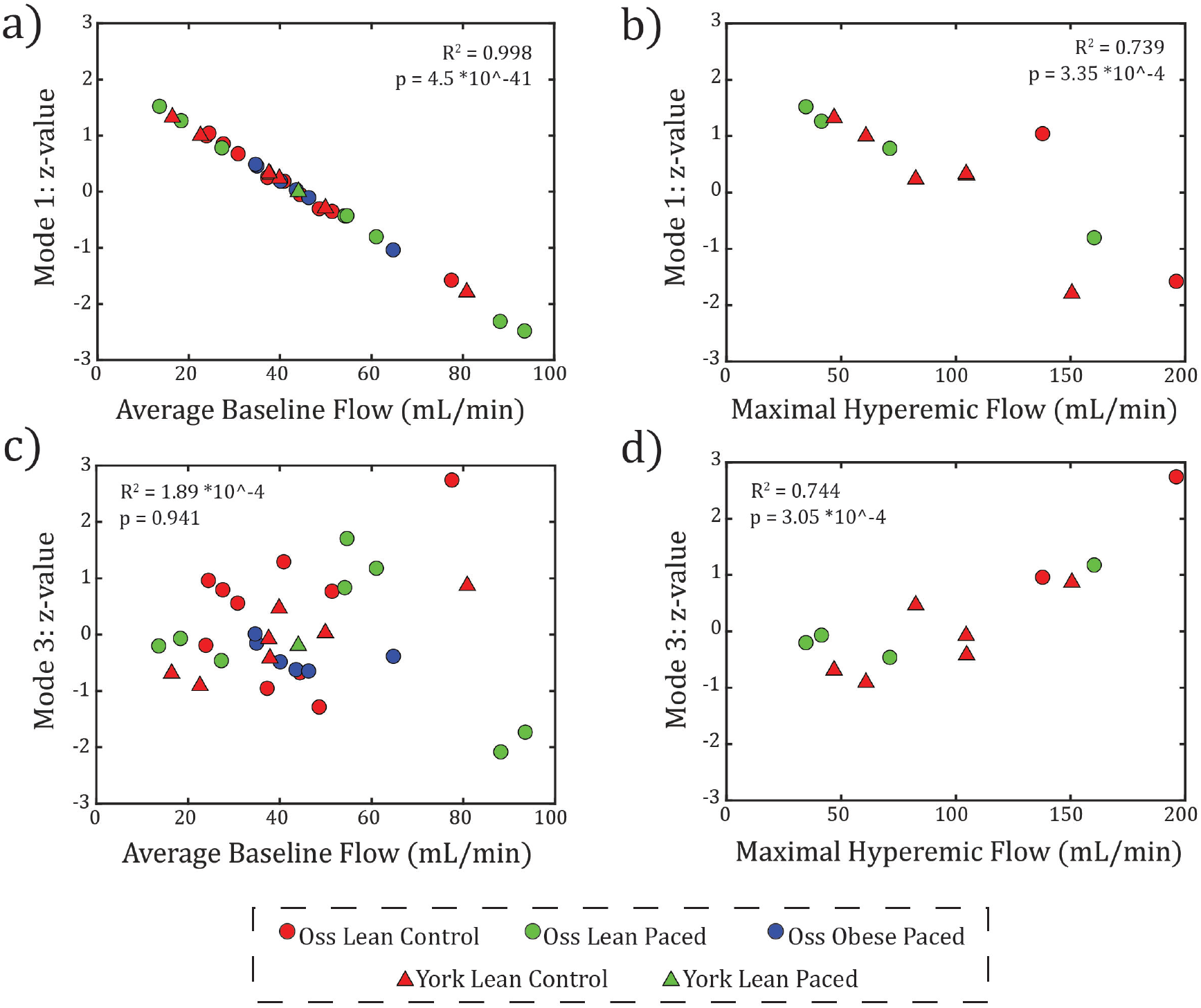
Correlations between SVD modes 1 and 3 and experimental data metrics of average baseline flow and maximum hyperemic flow.

To understand how SVD modes 1 and 3 correlate with maximum hyperemic flow, one- and two-variable linear regression models were constructed. Scatter plots of predicted hyperemic flow to measured values is shown in Figure S2. Table 1 displays the *R*^2^ and MSE values, measuring the explained variance and average magnitude of predicted errors, respectively. One-variable linear regression models of predicting hyperemic flow from average baseline flow was also used as a comparison. The two-variable regression model using the first and third SVD modes performed better than all single variable regression models.

**Table 1:**
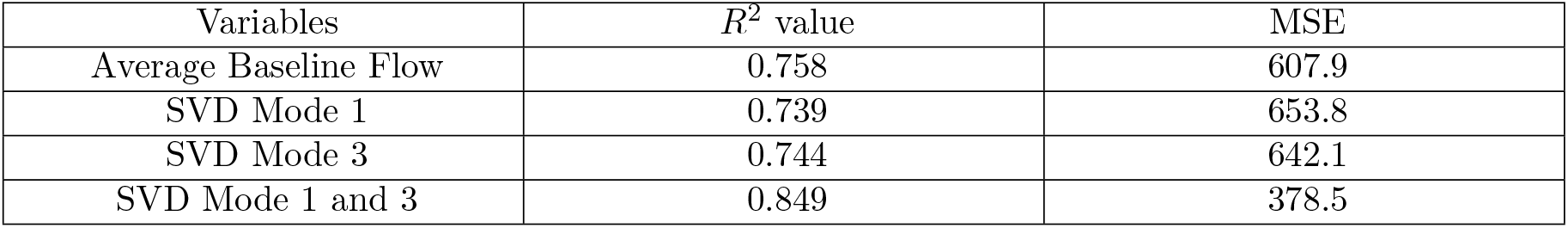
Linear Regressions for Estimating Hyperemic Flow.

### 3.4 Representative Waveforms for Experimental Groups

Waveforms constructed using average features from Ossabaw pigs in three experimental groups (*a*_*avg*_(*LeanControl*), *a*_*avg*_(*LeanPaced*), and *a*_*avg*_(*ObesePaced*)) are shown in Figure 8a. Panels b, c, and d of Figure 8 show the modal contributions of select SVD modes for each experimental group. In these panels, the contribution of the SVD modes(s) of interest were set by experimental group means for the Ossabaw pigs (*µ*(*LeanControl*), *µ*(*LeanPaced*), and *µ*(*ObesePaced*)) and the contribution of the remaining SVD modes was set by the modal average contribution across all pigs *µ*. Experimental group average and standard deviations for z-scores and selected experimental data (average coronary flow, heart rate, MBP, and LVEDP) are included in Table 2.

**Table 2:** Ossabaw experimental group SVD z-scores and hemodynamic data.

| Exp. Group | $z_1$ value | $z_2$ value | $z_3$ value | $z_4$ value | $z_5$ value | Avg Flow (mL/min) | Heart Rate (BPM) | MBP (mmHg) | LVEDP (mmHg) |
| --- | --- | --- | --- | --- | --- | --- | --- | --- | --- |
| Lean Control | $0.17 \pm 0.76$ | $-0.06 \pm 1.04$ | $0.40 \pm 1.14$ | $0.52 \pm 0.76$ | $-0.26 \pm 0.92$ | $40.6 \pm 16.2$ | $65.7 \pm 19.5$ | $126 \pm 37$ | $12.0 \pm 8.3$ |
| Lean Paced | $-0.36 \pm 1.41$ | $-0.20 \pm 1.13$ | $-0.10 \pm 1.25$ | $-0.44 \pm 1.29$ | $-0.22 \pm 0.64$ | $51.3 \pm 30.1$ | $75.1 \pm 17.8$ | $112 \pm 27$ | $22.1 \pm 12.5$ |
| Obese Paced | $0.01 \pm 0.51$ | $-0.37 \pm 0.55$ | $-0.38 \pm 0.24$ | $-0.40 \pm 0.50^*$ | $0.69 \pm 0.69^\dagger$ | $44.0 \pm 11.1$ | $64.0 \pm 8.9$ | $107 \pm 12$ | $23.9 \pm 4.37^*$ |

Note. Values are reported as mean ± standard deviation.
\* indicates $p < 0.05$ compared with Lean Control.
$^\dagger$ indicates $p < 0.05$ compared with Lean Paced.

**Figure 8:**
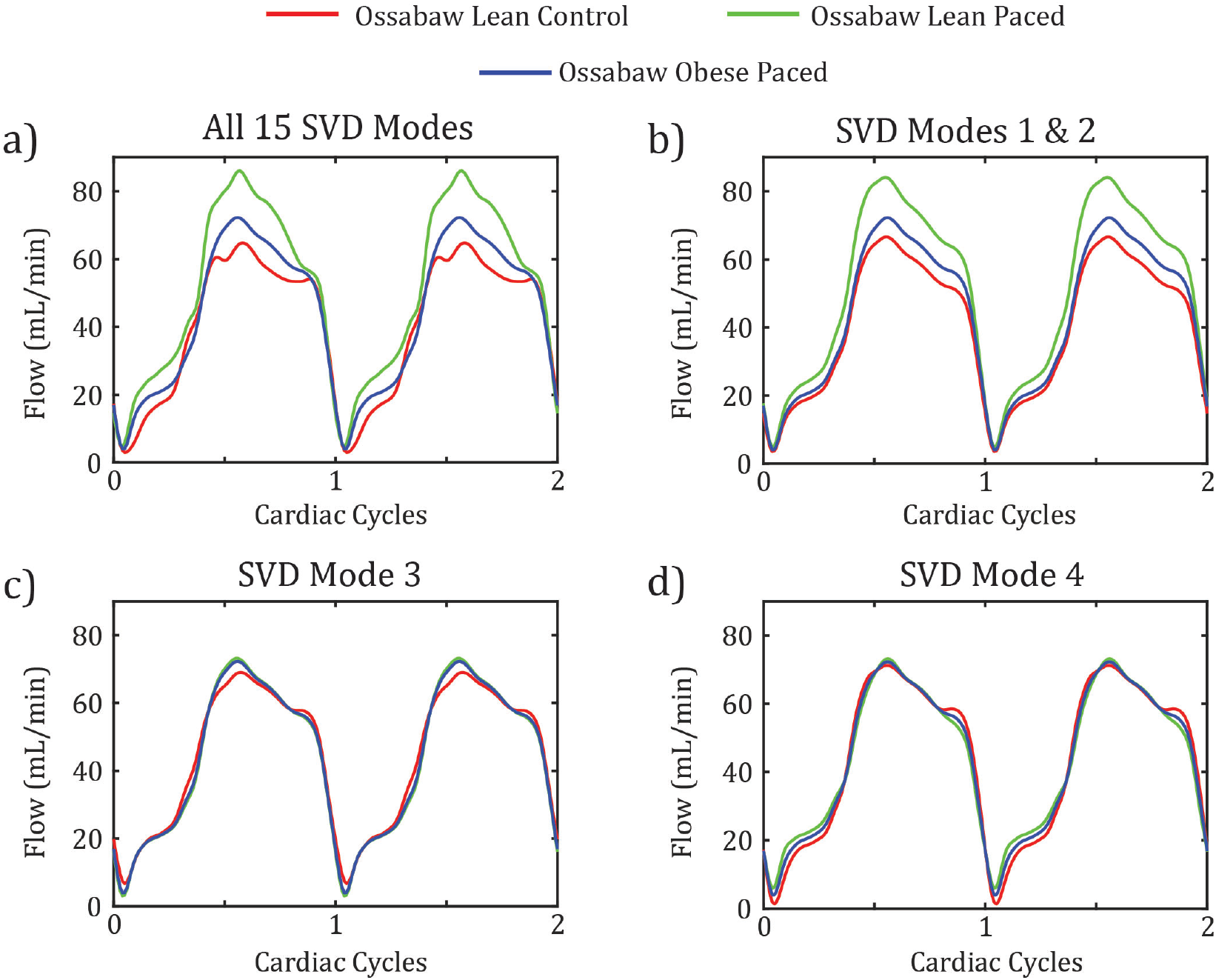
Comparison of reconstructed waveforms and modal contributions for Ossabaw pigs in the three experimental groups. Panel a shows the average waveforms for each group constructed from all 15 SVD modes. Panels b, c, and d show the contribution of select SVD modes (1 & 2, 3, and 4, respectively) for each experimental group. In each panel b-d, the modal contributions of the select SVD modes represent the average for each experimental group and the remaining modal contributions reflect the average across all pigs *a*_*avg*_. All waveforms begin in early systole and include two complete cardiac cycles.

Contributions of the first and second modes (Figure 8b) account for the differences in the magnitude and phase shift of the representative experimental group waveforms (Figure 8a). The trend in average z-score for SVD mode 1 (*z*_1_) matches the trend in average baseline flow between the three experimental groups, with average *z*_1_ decreasing as average baseline flow increases (Table 2). Flow waveforms constructed with only the average experimental-group-contribution for the third (Figure 8c) and fourth (Figure 8d) SVD modes show small differences between the control and paced groups that are otherwise obscured in the representative waveforms constructed from all SVD modes (Figure 8a). Trends in average z-scores for SVD modes 3 and 4 reflect trends in MBP and LVEDP for each experimental group (Table 2).

## 4 Discussion

This study is motivated by the need to develop a quantitative description and analysis of temporal patterns in LAD flow waveforms. Dimension reduction is used to simplify these waveforms into a small and uniform number of features. Recent work by Geddes and Randles proposed a methodology in which the coronary flow waveforms were characterized by various points-of-interest throughout the cardiac cycle, using synthetically generated waveforms [9]. In this study, we apply a different approach in which each coronary flow waveform is represented by a series of sinusoidal functions at harmonics of the cardiac cycle frequency. Therefore, our method enables reconstruction of coronary flow waveforms from a given set of features of the original signal (Equation 5). Errors associated with the reconstruction were identified based on the MSE and NMSE between the original waveform and reconstructed waveforms. MSE and NMSE values stabilized after approximately 10-15 harmonics, with further harmonics providing little useful information. The need for information from only the first 10-15 harmonics of the cardiac cycle would allow this technique to be applied to clinical data with low sampling frequencies. Samples with high MSE often showed high variability in cardiac cycle length (CCV), had associated spectral bleeding, or displayed changes in coronary flow waveform shape over the 30 cardiac cycles.

In the SVD analysis, the first mode accounted for most of the variance (Figure 4a). The large contribution of the first SVD-mode and near zero contribution of the remaining SVD modes (Figure 4b) indicates this mode predominantly defined the shape of the average flow waveform (black lines in Figure 5b) with diastolic dominant flow, a flow minimum during ventricular contraction, and a minor flow peak during mid-systole. The small percent variance explained by the remaining nodes indicates that the flow waveform of each pig only had minor variations from this diastolic-dominant pattern.

The second SVD mode had significant positive correlations with heart rate (Figure 5d). As heart rate increases, the amount of time the heart spends in its relaxed state decreases, and the systolic-to-diastolic time ratio increases [15]. Larger contributions of the second SVD mode increased the percent of the cardiac cycle with low coronary flow, and decreased the percent with high coronary flow (Figure 5c), representative of increases in the systolic-to-diastolic time ratio. In addition to the delay in the diastolic flow period, the second SVD mode also changed the shape of the diastolic flow peak. With negative contributions of the second mode (indicating lower heart rates), the diastolic flow peak becomes more skewed, and a larger percentage of the coronary flow occurs in early diastole. Davies et al. previously hypothesized that the sudden release of extravascular pressure during ventricular relaxation causes the peak in diastolic flow [8]. The effects of the sudden release of extravascular pressures compared to mid and late diastole are potentially more apparent at lower heart rates, when the heart spends more time in the relaxed state.

The third SVD mode had significant correlations with mean blood pressure and maximal hyperemic blood flow (Figure 6). Coronary stress tests were performed for a subset (12) of the 32 pigs included in this study. Positive correlation trends for hyperemic flow vs average resting blood flow and hyperemic blood flow vs MBP (not shown) were observed in the data. Interestingly, although the third SVD mode correlated with hyperemic flow, it did not correlate with average baseline flow (Figure 7). Additionally, multivariable linear regression with the first and third SVD modes provided better results than single-variable linear regression models using either average baseline flow or either SVD mode by itself (Table 1). This could suggest that the third SVD mode may encode additional information about microvascular function.

The fourth SVD mode primarily altered coronary flow during systole (Figure 5g). This mode did not present significant correlations with any hemodynamic quantity. However, non-significant correlations were observed with LVEDP (p=0.00268) (Figure 5h). Elevated filling pressures are a defining feature of heart failure [16]. Pigs within the pacing protocol were previously reported to have higher end-diastolic filling pressures compared to control (non-paced) pigs [13]. Therefore, this could indicate that the fourth SVD mode could be used as an indicator of heart failure.

Furthermore, the fourth SVD mode also has a non-significant positive correlation with Endo-Epi values (p=0.0873) (Figure S1). Computational models suggest that systolic coronary flow preferentially perfuses the subepicardium due to the spatial variation of intramyocardial pressures [3]. While this rationale supports the hypothesis that alterations to systolic flow with the fourth SVD mode would correlate with endocardial-to-epicardial flow ratios, further research would be required to support these claims.

The dimension reduction methods presented here demonstrate the ability to study *in vivo* coronary flow waveforms with more rigor than has previously been performed. Variability in heart rate and average blood flow may obscure the effects of microvascular or cardiac disease on coronary flow waveforms. This method demonstrates the ability to decompose coronary flow waveforms and isolate these effects. Higher SVD modes may provide information on mechanical or structural differences between waveforms. Additionally, they may contain information useful for predicting other clinical measurements, such as maximal hyperemic blood flow. Furthermore, these methods can be extended beyond *in vivo* waveforms and could be used to analyze or generate synthetic waveforms. Synthetic coronary flow waveforms can be projected into SVD spaces defined by *in vivo* data, potentially assisting with parameter estimation or sensitivity. Additionally, for computational models that rely on coronary flow waveforms as inputs, low-rank approximations from these SVD results could be useful for developing synthetic waveforms with physiological insight. For example, synthetic waveforms could be constructed from the first four SVD modes, with estimates of average coronary flow, heart rate, mean blood pressure, and LVEDP.

## 5 Limitations and Future Work

This study included a relatively low sample size of 32 pigs. Two different breeds of pigs (Yorkshire and Ossabaw) and three different experimental conditions (lean control, lean paced, and obese paced) were included in this study in an effort to increase sample size and perform a more generalizable analysis of coronary flow waveforms in heart and disease. However, the variability of coronary flow waveforms within this study likely does not cover the variability of data seen clinically.

In addition to low sample size, limited variability and uncertainty of experimental data metrics hinder the correlation analysis. For example, coronary stress tests were performed for only 12 of the 32 pigs. Clinically, patients with CFR measurements below 2 are categorized as having microvascular disease, whereas healthy patients may have higher CFR values ranging from 3 to 5 [17]. In this dataset, CFR values for lean paced Ossabaw pigs and lean control Yorkshire pigs ranged from 1.86-2.78, and the only CFR value recorded for a lean control Ossabaw pig (5.64) appeared as an outlier in this dataset. Higher sample sizes and a wider distribution of CFR values would be required to identify correlations with SVD modes. Furthermore, Endo-Epi ratios of 1.1-1.5 are typically reported in the literature [18], and low ratios are not expected in healthy hearts under normal resting conditions. In the data used in this study, Endo-Epi ratios varied from 0.44 to 1.39, and presented no significant trends between the different experimental conditions.

Future work will investigate how similar methods may be applied to analyze patient data using larger clinical datasets with more information. For example, patient coronary flow waveforms can be measured using transit-time flow measurements (TTFM) [19]. Additionally, datasets with information collected from coronary stress tests or positron emission tomography (PET) would provide information regarding hyperemic flow states and regional flow differences, respectively.

## 6 Conclusion

Overall, this work presents a method for analyzing shape and temporal patterns of coronary flow waveforms using Fourier transforms and SVD. Simplifying coronary waveforms to a small number of features that represent sinusoidal components at harmonics of the cardiac cycle frequency allows for the comparison of coronary flow waveforms collected at different heart rates or with different sampling frequencies. Further, we demonstrate that SVD analysis using these reduced features can provide insights into temporal patterns that exist in the coronary flow waveforms. While clinical and experimental analysis of coronary hemodynamics typically focuses on average values, the analysis presented in this study demonstrates the possibility of gaining additional physiological insights by studying temporal coronary flow patterns.

## Acknowledgments

This study is supported by NIH grants R01-HL158723 (DAB, CAF, JDT), T32-GM150581 (VES) and F31 HL179995 (VES).

## Ethics Approval Statement

All the protocols involving animals conformed to the National Institutes of Health Guide for the Care and Use of Laboratory Animals and were approved by the University of North Texas Health Science Center Institutional Animal Care and Use Committee

