## Supplemental Figures for "Shape Analysis of Coronary Flow Waveforms using Singular Value Decomposition"

### Supplementary Material

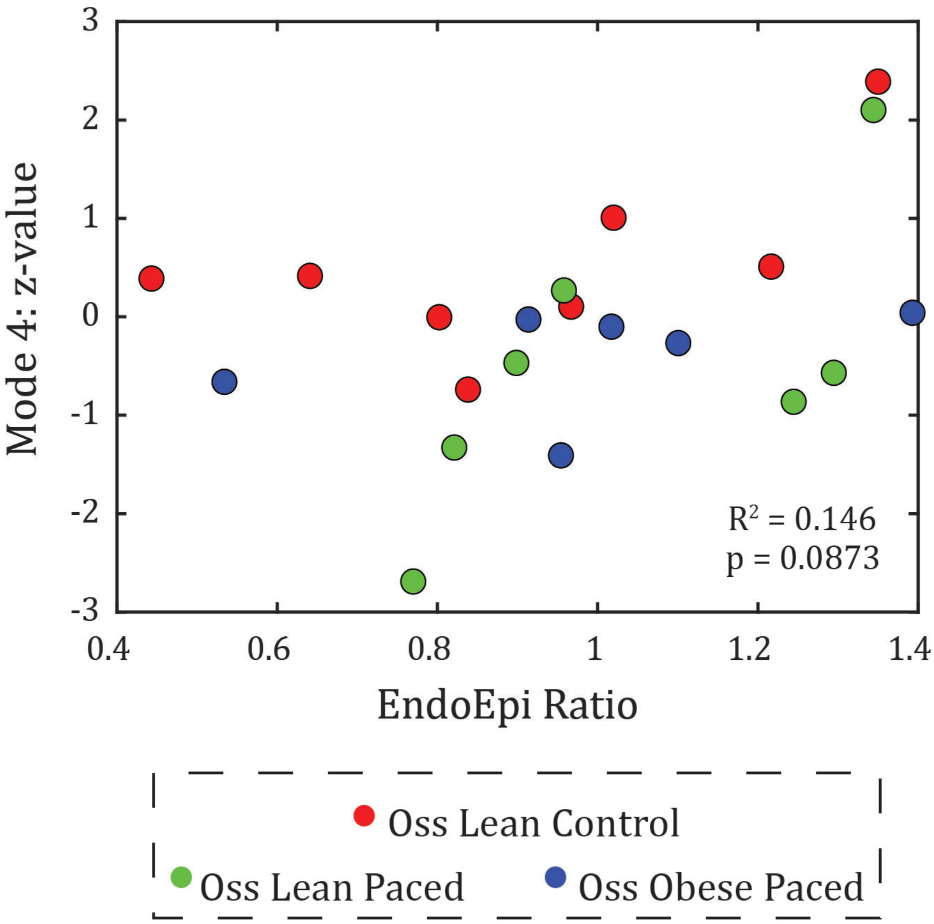


Figure S1: Correlation between SVD mode 4 and Endo-Epi ratios. Individual pigs are labeled by their experimental group: Ossabaw Lean Control (red), Ossabaw Lean Paced (green), Ossabaw Obese Paced (blue). No Endo-Epi data was collected for Yorkshire pigs.

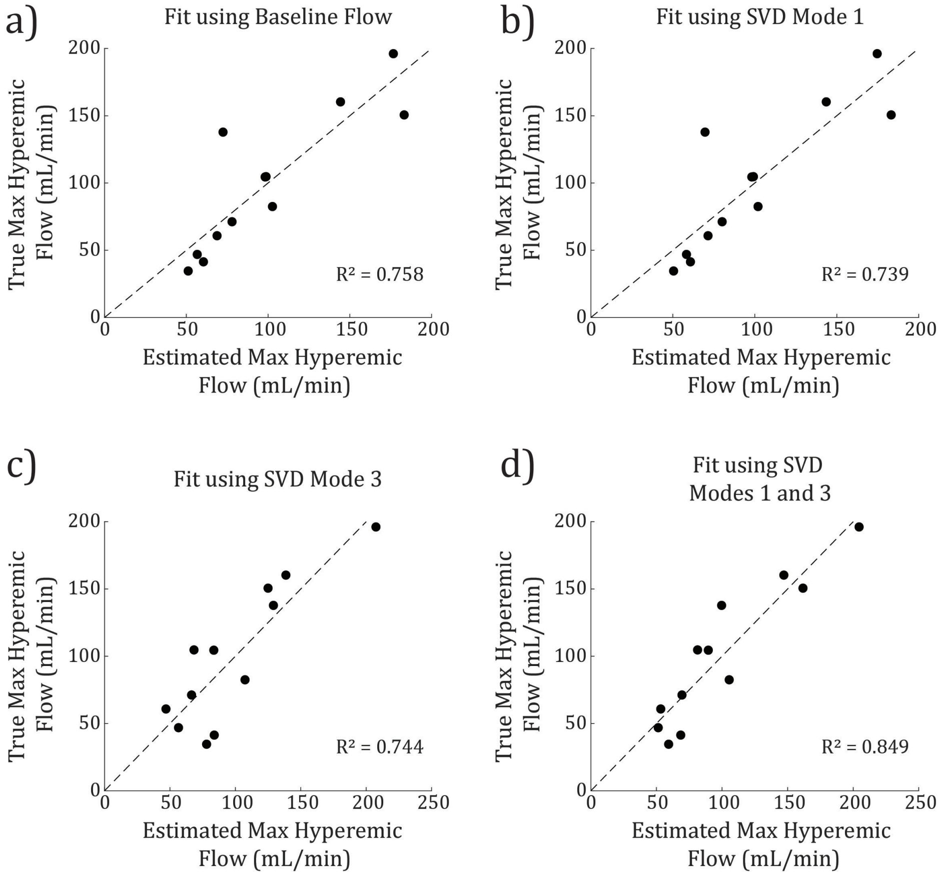


Figure S2: Comparisons between true hyperemic flow measurements and hyperemic flow measurements predicted by fitting regression lines to (a) average baseline flow measurements, (b) right singular values from SVD mode 1, (c) right singular values from SVD mode 3, and (d) right singular values from SVD modes 1 and 3
